# Dual Roles of the Retinal Pigment Epithelium and Lens in Cavefish Eye Degeneration

**DOI:** 10.1101/830885

**Authors:** Li Ma, Mandy Ng, Corine M. van der Weele, Masato Yoshizawa, William R. Jeffery

## Abstract

*Astyanax mexicanus* consists of two forms, a sighted surface dwelling form (surface fish) and a blind cave-dwelling form (cavefish). Embryonic eyes are initially formed in cavefish but they are subsequently arrested in growth and degenerate during larval development. Previous lens transplantation studies have shown that the lens plays a central role in cavefish eye loss. However, several lines of evidence suggest that additional factors, such as the retinal pigment epithelium (RPE), which is morphologically altered in cavefish, could also be involved in the eye regression process. To explore the role of the RPE in cavefish eye degeneration, we generated an albino eyed (AE) strain by artificial selection for hybrid individuals with large eyes and a depigmented RPE. The AE strain exhibited an RPE lacking pigment granules and showed reduced expression of the RPE specific enzyme retinol isomerase, allowing eye development to be studied by lens ablation in an RPE background resembling cavefish. We found that lens ablation in the AE strain had stronger negative effects on eye growth than in surface fish, suggesting that an intact RPE is required for normal eye development. We also found that the AE strain develops a cartilaginous sclera lacking boney ossicles, a trait similar to cavefish. Extrapolation of the results to cavefish suggests that the RPE and lens have dual roles in eye degeneration, and that deficiencies in the RPE may be associated with evolutionary changes in scleral ossification.

## Introduction

Closely related species or different forms of the same species offer unique opportunities to study the evolution of development (Brakefield, French & Zwann et al., 2003; Goldstein, 2010; Jeffery, 2016). A prime example is the teleost *Astyanax mexicanus*, which consists of two forms with strikingly different phenotypes (Jeffery, 2001; Gross, 2012; Keene, Yoshizawa, & McGaugh, 2015). The surface-dwelling form (surface fish) has large eyes and abundant melanophores, whereas cave-dwelling forms (cavefish), which consist of about 30 distinct populations (Gross, 2012), have reduced or lost both of these traits. Many other differences have evolved between surface fish and cavefish since their initial divergence from common ancestors about 20,000-200,000 years ago (Fumey, Hinaux, Noirot, Thermes, Rétaux et al., 2018; Herman, Brandvain, Weagley, Jeffery, Keene et al, 2018), including changes in skeletal structures (Yamamoto, Espinasa, Stock & Jeffery, 2003; Dufton, Hall & Franz-Odendaal, 2012; Gross, Krutzler & Carlson, 2014; Atukorala & Franz-Odendaal, 2018), olfactory, taste, and lateral line organs (Blin, Tine, Meister, Elipot, Bibliowicz et al., 2018; Yamamoto, Byerly, Jackman, & Jeffery, 2009; Teyke, 1990; Yoshizawa, Jeffery, van Netten, & McHenry, 2014), and behaviors (Yoshizawa, 2015). An important asset of the *A. mexicanus* system is that interbreeding offers a powerful genetic tool for interrogating the developmental mechanisms of evolutionary change (Borowsky & Wilkens; 2001; Protas, Conrad, Gross, Tabin, & Borowsky 2007; O’Quin, Yoshizawa, Doshi, & Jeffery, 2013). Accordingly, genetic crosses and molecular analysis have confirmed that a single gene (*oca2*) is involved in cavefish pigment loss (Sadoglu, 1957; Protas, Hersey, Kochanek, Zhou, Wilkens et al., 2006; Klaassen, Wang, Adamski, Rohner, & Kowalko, 2018), whereas about 12 different genes, each controlling a small part of the cavefish optic phenotype, are involved in eye loss (Borowsky & Wilkens, 2001; Protas et al., 2007; O’Quin et al., 2013).

One of the most dramatic phenotypic changes in cavefish is the absence of eyes (Krishnan & Rohner, 2017). Eyes are lacking or present as reduced structures in the adults of most *Astyanax* cavefish populations. It is known that cavefish embryos initiate eye formation, however, and the embryonic eyes are subsequently lost by a series of regressive events during larval development (Cahn, 1958; Wilkens, 1988; Jeffery, 2009). First, a smaller neuro-ectodermal eye field and optic vesicle are formed in cavefish compared to surface fish embryos (Yamamoto, Stock, & Jeffery et al., 2004). Second, after optic cup formation and lens induction, a slightly smaller presumptive lens and retina, the latter with a reduced ventral portion, develop in cavefish embryos (Alunni, Menuet, Candal, Penigault, Jeffery et al., 2007). Third, about a day after hatching, the lens undergoes massive apoptosis, apoptotic cell death then spreads to the retina, which becomes disorganized, photoreceptor cells are initially formed but then ultimately degenerate, and eye growth is slowed and eventually ceases (Jeffery and Martasian, 1998; Strickler, Yamamoto & Jeffery, 2007; Jeffery, 2009). Accessory optic structures, such as the cornea, iris, and sclera, either fail to develop or only partially differentiate in cavefish (Strickler. Yamamoto & Jeffery, 2001). For example, the small cavefish sclera is highlighted by a cartilage ring but is generally lacking the two boney ossicles typical of the surface fish sclera (Yamamoto et al., 2003; O’Quin, et al., 2013; O’ Quin, Doshi, Lyon, Hoenemeyer, Yoshizawa, et al., 2017). As the cavefish larva grows, the small degenerating eyes gradually sink into the orbits and are covered by skin and connective tissue.

Many of the changes involved in cavefish eye degeneration have been attributed to apoptosis and subsequent dysfunction of the lens. When a surface fish embryonic lens was transplanted into a cavefish optic cup, after its own degenerating lens was removed, the surface fish lens remained functional in the cavefish host, overall eye growth was rescued, and the iris, cornea, an organized layered retina with photoreceptor cells, and an ossified sclera were formed, suggesting that the lens plays a major role in cavefish eye degeneration (Yamamoto and Jeffery, 2000). The importance of the lens in organizing corneal and retinal development has also been demonstrated in zebrafish and other vertebrates (Coulombre & Coulombre, 1964; Breitman, Clapoff, Rossant, Tsui, Glode et al., 1987; Kaur, Key, Stick, McNeisch, Akeson et al., 1989; Kurita, Sagara, H., Aoki, Y., Link, B. A; Beebe & Coates, 2000; Asherley-Padan, Marquart, Zhou, & Gruss, 2000; Pandit, Jidigam, Patthey & Gunhaga, 2015). However, cavefish eyes rescued by lens transplantation fail to attain the large size of typical surface fish eyes (Yamamoto & Jeffery, 2000; Strickler, et al. 2007) and, at least in some cases, vision does not appear to be restored (Romero, Green, Romero, Lelonek, & Stropnicky, 2003). These results suggest that the lens, although important, is not the only factor involved in cavefish eye degeneration, and that other optic elements, or even factors operating outside the eyes, may also be involved. The existence of multiple causes of cavefish eye degeneration is further supported by lens extirpation in surface fish embryos, which results in smaller but morphologically normal retinas in adults, contrary to the complete eye loss expected if the lens was the only factor involved in eye degeneration (Strickler et al., 2007). Finally, as described above, genetic crosses between surface fish and cavefish indicate that eye degeneration is a complex trait controlled by multiple genes that could function in different parts of the developing eyes.

Strickler et al. (2007) proposed the dual signal model to address the complexity of cavefish eye degeneration. The dual signal model hypothesizes a requirement for two or more individual optic elements, which secrete signals that together are necessary for complete eye development and growth in *A. mexicanus*. According to the model, all of the required optic elements and signals would be present in surface fish, resulting in full eye development, whereas all of these elements and signals would be missing or modified in cavefish, resulting in eye loss. Thus, when the lens is removed in surface fish, only one of these elements and signals would be lost, resulting in a smaller eye. And when a lens is transplanted from surface fish into cavefish only one of the missing elements/signals, the lens, would be replaced, producing a complete but small eye. Strickler et al. (2007) further proposed that the retinal pigment epithelium (RPE) may be another optic structure that is involved in cavefish eye degeneration.

The RPE is a monolayer of melanin pigmented cells that performs many critical visual functions, including prevention of light scatter by absorbing extraneous photons, conversion of all-trans-retinol to 11-cis-retinol in the visual cycle, and nourishment and protection of the underlying photoreceptor layer (Strauss, 2005). The possibility that the RPE is defective in cavefish is supported by the absence of melanization in larval Pachón cavefish eyes (McCauley, Hixon, & Jeffery, 2004; Bilandzija, Ma, Parkhurst & Jeffery, 2013) and less extensive eye degeneration in those cavefish populations with a hypopigmented RPE, such as Tinaja and Chica (Zilles, Tillmann, & Bennemann, 1983). Furthermore, molecular ablation experiments in other vertebrates have shown that the RPE is essential for normal eye development (Raymond & Jackson, 1995; Rymer & Wildsoet, 2005; Hanovice, Leach, Slater, Gabriel, Romanovicz et al., 2019). Therefore, it is important to determine the role of the RPE in *A. mexicanus* eye development in order to fully understand the mechanisms of cavefish eye degeneration.

In the present study, we have examined the role of the RPE in eye development and cavefish eye degeneration in an *A. mexicanus* albino eyed (AE) strain constructed by artificial selection. We showed that the AE strain has a defective RPE and found that lens ablation in the AE strain has stronger effects on eye development than in surface fish. The results support a dual role for the RPE and lens in cavefish eye degeneration as well as an impact of the defective RPE on cavefish scleral differentiation.

## Materials and Methods

### Biological materials and husbandry

*Asytanax mexicanus* surface fish and cavefish were obtained from Jeffery laboratory stocks. The surface fish laboratory stocks were originally collected at Balmorhea State Park, TX, USA and Nacimiento del Rio Choy, Tamaulipas, Mexico, and Pachón cavefish laboratory stocks were originally obtained in Cueva de El Pachón, Tamaulipas, Mexico. Fish were raised in the laboratory at 25°C on a 14-hr light and 10-hr dark photoperiod (Jeffery, Strickler, Guiney, Heyser, and Tomarev, 2000). Embryos were obtained by natural spawning or by *in vitro* fertilization (see below) and raised at 23-25°C. Larvae were fed brine shrimp beginning at 5-6 days post-fertilization (dpf), and adults were fed TetraMin Pro flakes (Tetra Holding Inc, Blacksburg VA). Animals were maintained and handled according to Institutional Animal Care guidelines of the University of Maryland, College Park (IACUC #R-NOV-18-59) (Project 1241065-1).

### Hybridization and artificial selection

The initial hybridization of P0 surface fish and cavefish and intercrossing of F1, F2, and F3 hybrids to form the AE strain was done by *in vitro* fertilization (Ma, Strickler, Parkhurst, Yoshizawa, Shi, et al., 2018). Unfertilized eggs were collected by applying gentle pressure to the abdomen of a gravid female. Sperm were obtained by gently squeezing the area around the cloaca of males. *In vitro* fertilization was carried out by mixing sperm with unfertilized eggs followed by dilution in fish system water. Adult hybrid males and females with large eyes and lacking melanin pigmentation were identified by visual inspection and crossed following the selection regime described in Figure 1.

**Figure 1.**
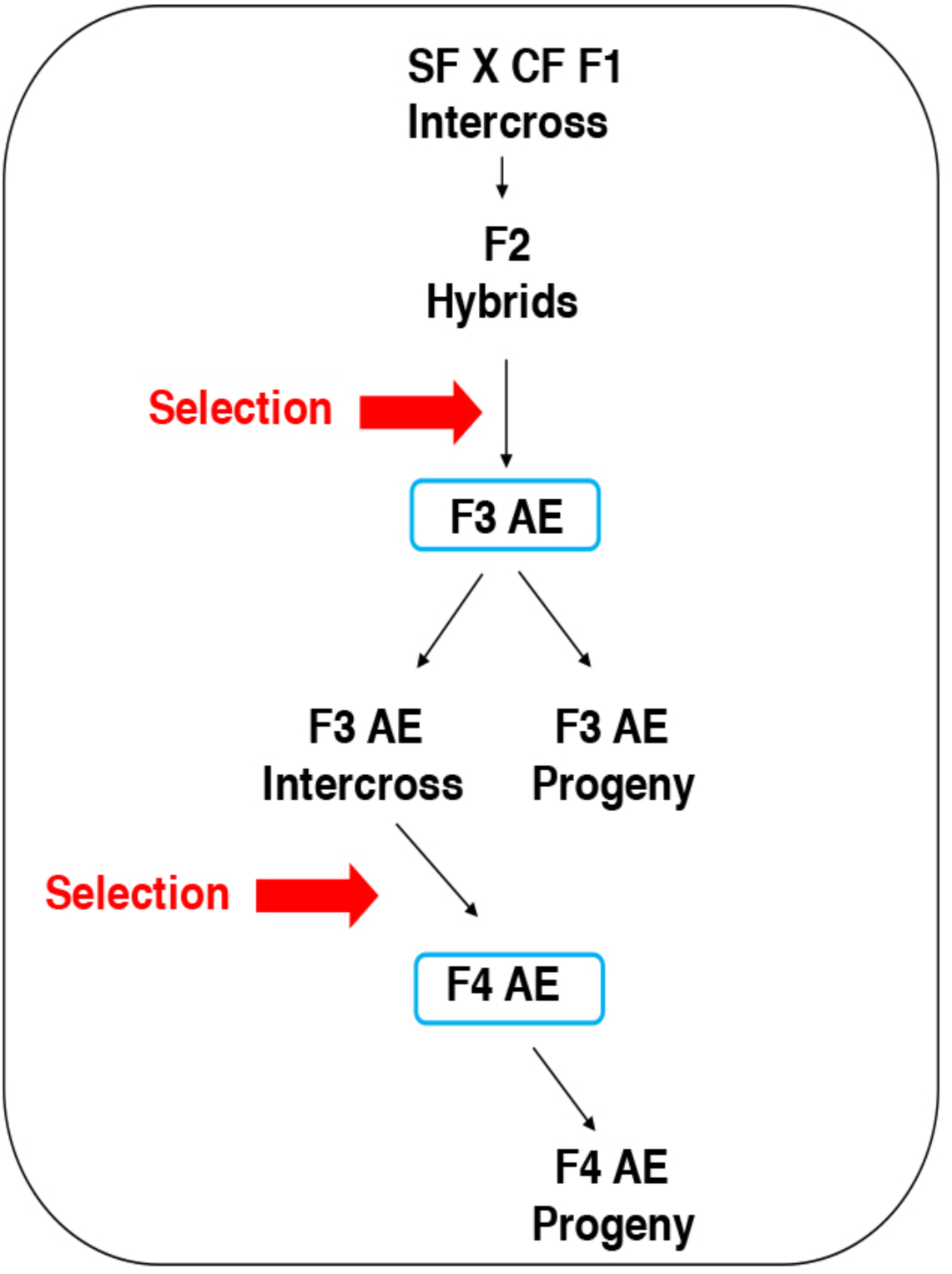
Flow chart for artificial selection. The hybridization and selection regime used to generate the albino eyed (AE) strain. See Materials and Methods and Results for details.

### Pigmentation analysis

Surface fish and AE adults were anesthetized with tricaine (2 μg/ml; Western Chemical Inc., Ferndale, WA) and examined for eye and body pigmentation while viewed under a stereomicroscope. Larval fin pigmentation was examined by removing a part of the dorsal tail fin with small dissection scissors. The pieces of tail fin were mounted under coverslips on glass microscope slides, and pigment cells were identified under a compound microscope.

### Lensectomy

The lens was removed from the optic cup on one side of surface fish and AE larvae at about 30 hr post-fertilization (hpf) by microdissection with Tungsten needles (Yamamoto & Jeffery, 2002). The operated eyes were examined by light microscopy, and larvae in which the optic cup was malformed by surgery were discarded. The operated larvae were cultured to the late larval and adult stages as described above.

### Eye measurements

Surface fish and AE larvae, fry, and adults were anesthetized with 2 μg/ml tricaine, and eyeball and pupil diameters were measured along the anterior posterior optic axis using a compound microscope equipped with Image J or AxioVision software (Carl Zeiss Microscopy GmbH, Jena, Germany) and photographed.

### Histology

Adult surface fish and AE were fixed in 4% paraformaldehyhde (PFA) in phosphate buffered saline (PBS) overnight at 4°C. The fixed specimens were washed in PBS, dehydrated in an increasing series of ethanol solutions up to 100% ethanol, and embedded in Paraplast (Polyscience, Inc, Worthington, NY). The blocks were sectioned at 8 μm, the sections were attached to glass slides, and the slides were stained with Harris hematoxylin-eosin.

### Immunostaining

Surface fish and AE larvae were fixed in 4% PFA overnight at 4°C, dehydrated through an increasing series of methanol solutions to 100% methanol, and stored at -20°C. The larvae were rehydrated, blocked with 2% blocking reagent (Roche, Basel, Switzerland), 2% bovine serum albumen, 5 or 10% heat inactivated goat serum in 100 mmol l^-1^ maleic acid, pH 7.5, and 150 mmol l^-1^ NaCl, and incubated in primary antibody. The zpr2 monoclonal antibody (Zebrafish International Resource Center, Eugene, OR) was used at 1:200 (F3 AE progeny and corresponding surface fish) or 1:500 dilutions (F4 AE progeny and corresponding surface fish). The rabbit rpe65 polyclonal antibody (Abcam, Cambridge, MA) was used at a 1:500 dilution for the AE strain and surface fish. The primary antibody incubations were carried out for 48 hr at 4°C, and the specimens were rinsed with superblock. Incubation with secondary antibody was carried out overnight at 4°C, the specimens were successively rinsed with PBS, 0.1% Triton X-100, and 0.2% BSA, and visualized by fluorescence microscopy. The secondary antibody for zpr2 staining was goat anti-mouse IgG Alexa Flour 488 (Therm Fisher, Waltham, MA) used at an 1:200 dilution. The secondary antibody for rpe65 staining was goat anti-rabbit IgG Alexa Fluor 488 (Invitrogen, Carlsbad, CA) used at a 1:200 dilution.

### Cartilage and bone staining

Surface fish, cavefish, and AE strain adults of the same size and age were stained for cartilage and bone using Alcian blue and Alizarin Red (Hanken & Wassersug, 1981) with the following modifications: 10% formalin fixation, Alcian blue staining, trypsin treatment, and alizarin red staining were each carried out at room temperature for 12 hr. Stained specimens were imaged by light microscopy and photographed.

### Statistical analysis

Statistical scores of eye and pupil sizes between F3 AE, F4 AE, and surface fish were determined by one-tailed Students t tests with p values and other relevant parameters described in Fig. S1. Statistical analysis of eye sizes in lens ablation experiments was done by three-way ANOVA (population, lensectomy and age) with post-hoc Bonferroni correction (Tables 1 and 2), and two-way ANOVA (population, age) with post-hoc Bonferroni correction (Tables 3 and 4). P values, F statistics, and other relevant parameters are described in Tables 1-4 and Tables S1 and S2. All statistical analyses were performed under SPSS statistics software (release 25.0.0.1; IBM, Armonk, NY) and Excel (Microsoft, Redmond, WA).

**Table 1.** Effect of Lens Deletion on Eye Size in Developing Progeny of F4 Albino Eyed (AE) Strain and Surface Fish (SF)

| Age (dpf) <sup>†</sup> | Type | Sample size (N) | Mean eye size at Control side (mm<sup>2</sup> ± s.e.m.) | Mean eye size at Lensectomy side (mm<sup>2</sup> ± s.e.m.) | p-value<sup>‡</sup> (Bonferroni correction) |
| --- | --- | --- | --- | --- | --- |
| 3 | SF | 36 | 337.9 ± 5.3 | 311.4 ± 7.5 | 0.0071 |
| 3 | AE | 36 | 314.0 ± 4.9 | 292.8 ± 8.8 | 0.0533 |
| 7 | SF | 28 | 521.6 ± 10.6 | 375.5 ± 8.1 | 2.45 × 10<sup>-5</sup> |
| 7 | AE | 32 | 375.5 ± 8.1 | 298.6 ± 10.4 | 2.45 × 10<sup>-6</sup> |
| 23 | SF | 14 | 1223.3 ± 114.6 | 647.2 ± 92.3 | 0.065 |
| 23 | AE | 24 | 721.2 ± 46.5 | 447.89 ± 32.9 | 8.61 × 10<sup>-5</sup> |
| 35 | SF | 10 | 11961.6 ± 41.7 | 761.2 ± 119.3 | 0.0093 |
| 35 | AE | 20 | 892.2 ± 52.3 | 415.9 ± 59.3 | 1.08 × 10<sup>-5</sup> |

<sup>†</sup>dpf (days post fertilization)
<sup>‡</sup>Significance determined by *t*-test between the eye sizes of lensectomy and control sides with Bonferroni correction.

**Table 2.** Statistical scores of three-way repeated ANOVA for Effect of Lens Deletion on Eye size

| Source | F-score | p-value |
| --- | --- | --- |
| Population* | F(1,92) = 163.3 | 3.56 × 10<sup>-75</sup> |
| Age* | F(3,92) = 184.6 | 4.20 × 10<sup>-22</sup> |
| Lensectomy* | F(1,92) = 304.7 | 6.00 × 10<sup>-31</sup> |
| Pop × Age* | F(3,92) = 32.7 | 1.81 × 10<sup>-14</sup> |
| Pop × Lens | F(1,92) = 0.4 | 0.539 |
| Age × Lens* | F(3,92) = 66.3 | 6.35 × 10<sup>-23</sup> |
| Pop × Age × Lens | F(3,92) = 0.4 | 0.766 |

\*: statistically significant (P < 0.05)

**Table 3.** Eye Size Ratio of Lensectomy and Control Eyes in Developing Progeny of F4 Albino Eyed Strain (AE) and Surface Fish (SF) dpf: days post-fertilization

| Age (dpf)<sup>†</sup> | Type<sup>‡</sup> | Eye Size Ratio<sup>§</sup> ± SEM | p-value<sup>¶</sup><br>(Bonferroni<br>correction) |
| --- | --- | --- | --- |
| 3 | SF | 0.918 ± 0.990 | 0.5961 |
| 3 | AE | 0.933 ± 0.027 |  |
| 7 | SF | 0.843 ± 1.621 | 0.2135 |
| 7 | AE | 0.798 ± 0.255 |  |
| 23 | SF | 0.768 ± 0.574 | 0.0327 |
| 23 | AE | 0.606 ± 0.058 |  |
| 35 | SF | 0.646 ± 0.646 | 0.0184 |
| 35 | AE | 0.356 ± 0.062 |  |

<sup>†</sup>dpf: days post-fertilization
<sup>‡</sup>Number of specimens is same as in Table S1.
<sup>§</sup>Ratio is mean eye size on lensectomy side/control side.
<sup>¶</sup>p value significance between SF and AE determined by *t*-test with post-hoc Bonferroni correction.

**Table 4.** Statistical scores of two-way ANOVA for Eye Size Ratio of Lensectomy

| Source | F-score | p-value |
| --- | --- | --- |
| Population* | F(1,92) = 15.4 | $1.69 \times 10^{-4}$ |
| Age* | F(3,92) = 46.3 | $2.45 \times 10^{-18}$ |
| Pop × Age* | F(3,92) = 4.0 | 0.010 |

\*: statistically significant ( $P < 0.05$ )

## Results

### Production of an Astyanax Albino Eyed Strain by Artificial Selection

A flow diagram illustrating the steps in production of the AE strain is shown in Figure 1. To establish the AE strain, we capitalized on independent segregation of surface fish-like eye and cavefish-like pigmentation trait, as determined by eye and pupil measurements and melanophore visualization in the eyes, trunk, and tailfin, in the F2 progeny of a surface fish x cavefish F1 intercross. After raising the F2 hybrids to adulthood, the small number of adults lacking pigmentation with large eyes (O’Quin et al., 2013) were identified by visual inspection, and crossed to obtain an F3 generation (F3 AE). The F3 AE were raised to adulthood, and selection of offspring for large eyes and albinism was repeated to obtain an F4 generation (F4 AE) (Fig. 2A). An additional round of F4 AE intercrossing, eye and pigment trait determination, and selection as described above did not yield an F5 AE generation with significant differences compared to the F4 AE generation. Thus, the artificial selection regime was terminated at the F4 generation. We used the original F3 AE and/or F4 AE adults and their spawned larvae and fry, termed F3 AE and F4 AE progeny respectively (Fig. 1), in the following analyses.

**Figure 2.**
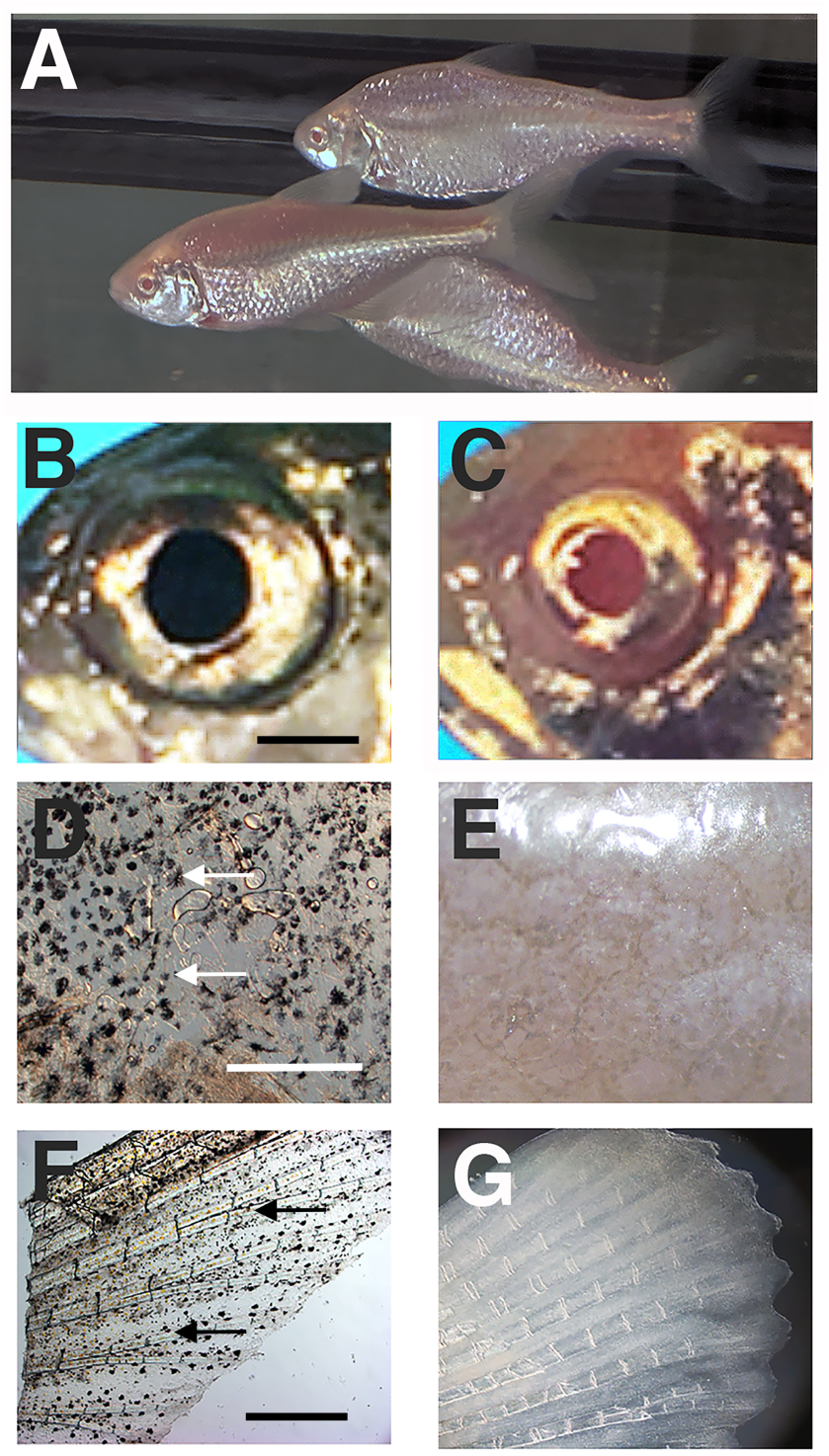
Characterization of eye and pigmentation phenotypes in the albino eyed (AE) strain. A. F4 AE adults. B, C. Eye comparisons. The pigmented eye of an adult surface fish showing melanin pigmented pupil (B) compared to the albino eye of an adult F4 AE with pupil lacking melanization (C). Scale bar in B: 50 mm; magnification is the same in B and C. D-G. Body pigmentation comparisons. Melanophores (arrows) are present in the trunk (D) and tail fin (E) of adult surface fish (D, F) but not the trunk (E) and tail fin (G) of F4 AE (E, G). Scale bar in D: 150 μm; magnification is the same in D and E. Scale bar in F: 400 μm: magnification is the same in F and G.

### Eye and Pigmentation Phenotypes in the Albino Eyed Strain

The eye and pigmentation phenotypes of the AE strain were compared to those of surface fish. Most of the F3 AE and all of F4 AE adults exhibited large eyes that were morphologically similar to surface fish eyes, although the mean eyeball and pupil (used as a proxy for the lens) sizes of both AE generations were slightly smaller than surface fish adults of comparable size and age (Fig. 2B, C, Fig. S1). The small differences in eye size between surface fish and the F3 and F4 AE strains were significant at p < 0.001 (Fig. S1). The AE adults lacked melanophores in the eyes, body, and fins (Fig. 2D-G). The results demonstrate that the cavefish albino and surface fish large eyed phenotypes can be merged in hybrids to form a novel *Astyanax* line via only a few rounds of artificial selection.

### Defective Retinal Pigment Epithelium in the Albino Eyed Strain

The absence of eye pigmentation suggested that the AE strain may be defective in RPE development. This possibility was explored by examining retinal differentiation in F3 AE and F4 AE larvae. We first compared the RPE in AE to surface fish of comparable size and age. In contrast to the highly pigmented RPE present in sections of surface fish retinas, the retinas of the AE strain showed RPEs without discernable melanin granules (Fig. 3A, B). Aside from differences in the RPE, retinal morphology was relatively normal in the AE strain, although the widths of retinal layers sometimes varied in different AE progeny compared to surface fish (Fig. 3A, B).

**Figure 3.**
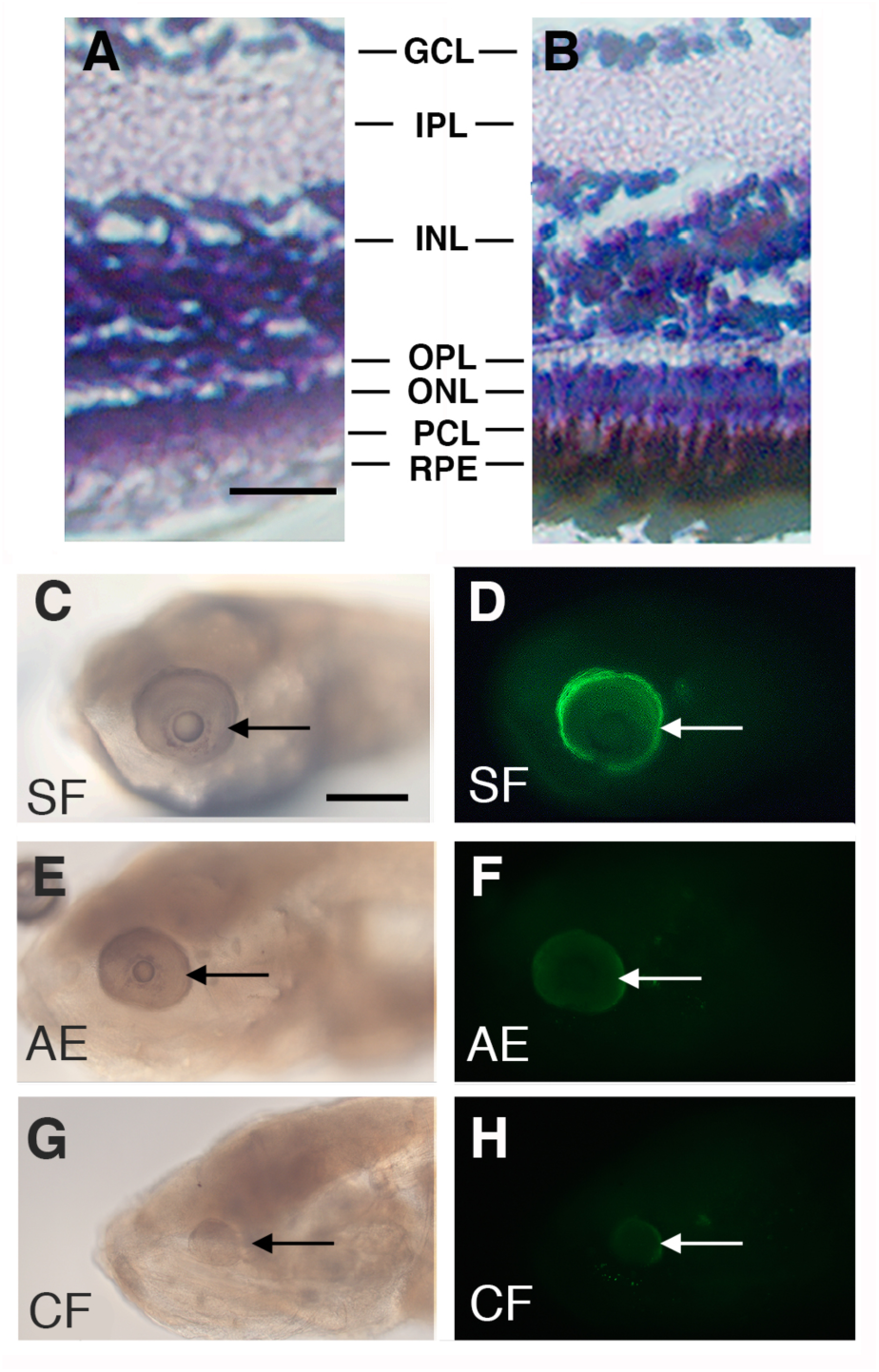
Defective retinal pigment epithelium (RPE) in the albino eyed (AE) strain. A-B. Retinal sections of adult surface fish and F4 AE stained with hematoxylin-eosin. The F4 AE (A) and surface fish (B) stained with hematoxylin-eosin at 14 days post fertilization (14 dpf). The RPE is lacking pigment in F4 AE surface fish but is melanized and interdigitates with the adjacent photoreceptor layer in surface fish. GCL: Ganglion cell layer. IPL: Inner plexiform layer. INL: inner nuclear layer. OPL: Outer plexiform layer. ONL: outer nuclear layer. PCL; Photoreceptor cell layer. RPE; Retinal pigment epithelium. Scale bar in A: 25 μm; magnification is the same in A and B. C-H. Immunostaining of 7 dpf larvae with rpe65 antibody. C-D. Surface fish. E-F. F4 AE strain. G-H. Pachón cavefish. The rpe65 antigen is brightly stained in surface fish eyes, but dim or not detectible in AE or cavefish eyes respectively. Arrows: eyes. Scale bar in C: 300 μm; magnification is the same in C-H.

To determine whether changes in addition to pigmentation occur in the AE RPE, surface fish, cavefish, and F4 AE larvae were stained with rpe65 antibody. The rpe65 antibody recognizes retinol isomerase, a critical enzyme in the vertebrate visual cycle that is specifically expressed in the RPE (Hamel, Tsilou, Harris, Pfeffer, Hooks, et al, 1993; Jin, Li, Moghrabi, Sun, & Travis, 2005). Robust expression of rpe65 antigen was observed in the eyes of all developing surface fish larvae, but rpe65 immunostaining was considerably reduced or undetectable in the eyes of all F4 AE and cavefish larvae, respectively (Fig. 3C-H). These results indicate that retinol isomerase expression is downregulated in the RPE of F4 AE larvae.

Both the RPE and photoreceptor layers show modifications in degenerating cavefish eyes (Yamamoto & Jeffery, 2000; Strickler & Jeffery, 2009). To determine whether the photoreceptor layer is affected in AE, we stained F3 AE and F4 AE progeny larvae with zpr2, an antibody that targets photoreceptor cell bodies in the zebrafish retina (Qin, Kidd III, Thomas, Poss, Hyde et al., 2011; Bader, Kusik, & Beharse, 2012). The retinas of most F3 AE larvae were stained with zpr2 (Fig. 4A-C), although a few F3 AE larvae, generally those with smaller eyes, lacked appreciable zpr2 staining (Fig. 4D-F). In contrast to F3 AE, zpr2 staining was present in the retinas of all tested F4 AE larva (Fig. 4G), implying that hybrids without a photoreceptor layer were likely purged from the F4 AE generation during artificial selection.

**Figure 4.**
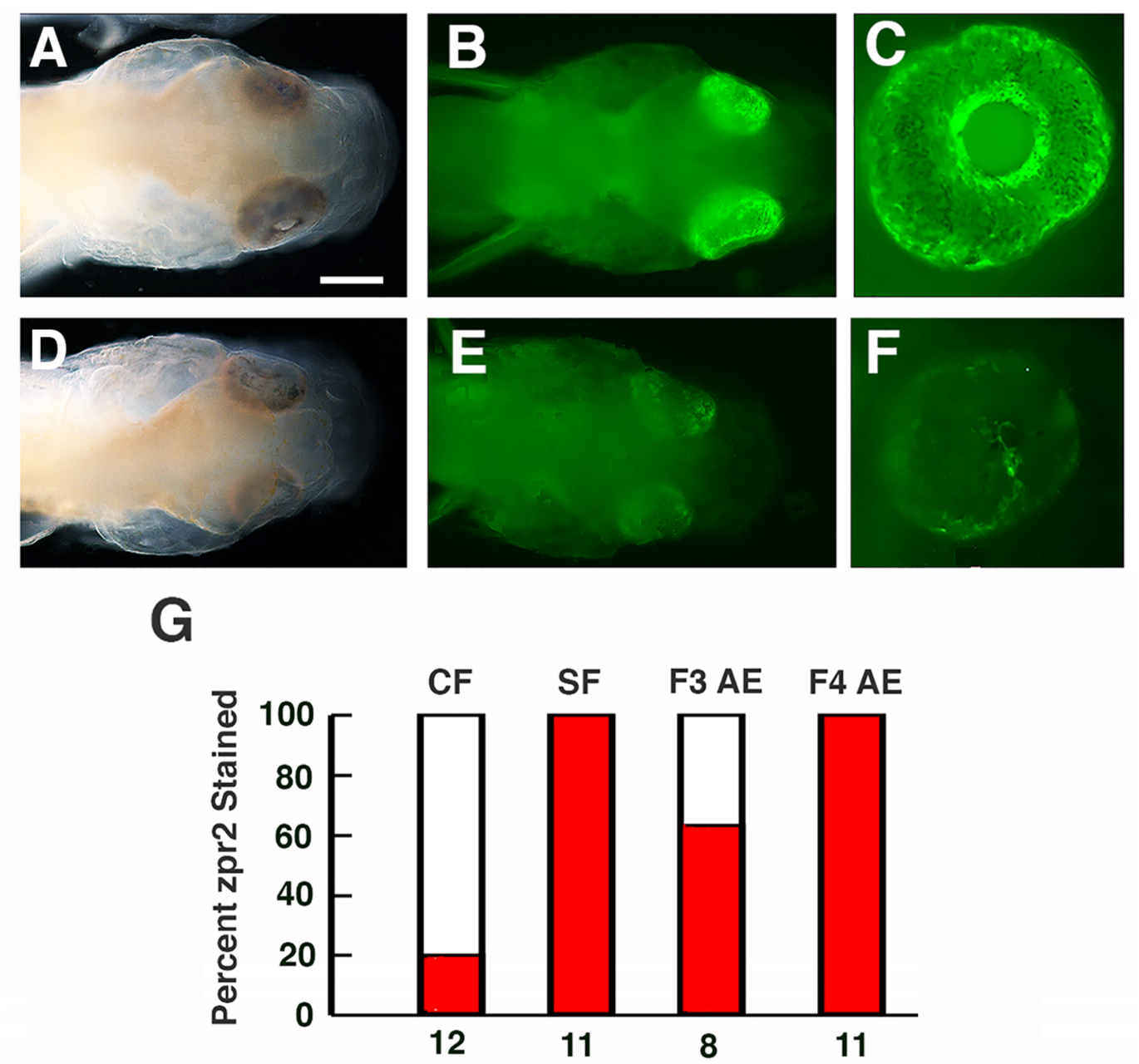
Retinal photoreceptor layer in the albino eyed (AE) strain determined by staining with zpr2 antibody at 3 days post fertilization. A-C. The majority of F3 AE larvae stain positively for zpr2 antigen. D-F. A minority of F3 AE larvae lack appreciable zpr2 staining. A, D. Bright field images. B, C and E, F. Fluorescence images. Scale bar in B: 300 μm; magnifications are the same in A-F. G. Bar graphs showing the percentage (red bars) of cavefish (CF), surface fish (SF), F3 AE, and F4 AE with eyes showing zpr2 staining. Number of samples assayed is shown at the bottom of each bar.

In summary, loss of melanin pigmentation and downregulation of retinoid isomerase indicated that RPE development is defective in the AE strain.

### Eye Size in the Albino Eyed Strain is Highly Sensitive to Lens Deletion

The AE strain offers the opportunity to determine the relative roles of the lens and RPE in regulating eye development by comparing the effects of lens deletion in AE and surface fish larvae. Accordingly, the lens vesicle was removed from the optic cup on one side of the head of F3 AE, F4 AE, and surface fish larvae at 30 hpf (Fig. 5 top). The lens vesicle was not disturbed on the opposite side of the head, which served as a control for the operated side. The AE and surface fish larvae were cultured for the next few weeks, and eyeball sizes were measured and compared on the operated and control sides of the head beginning at 3-day post fertilization (dpf) and periodically during subsequent development.

**Figure 5.**
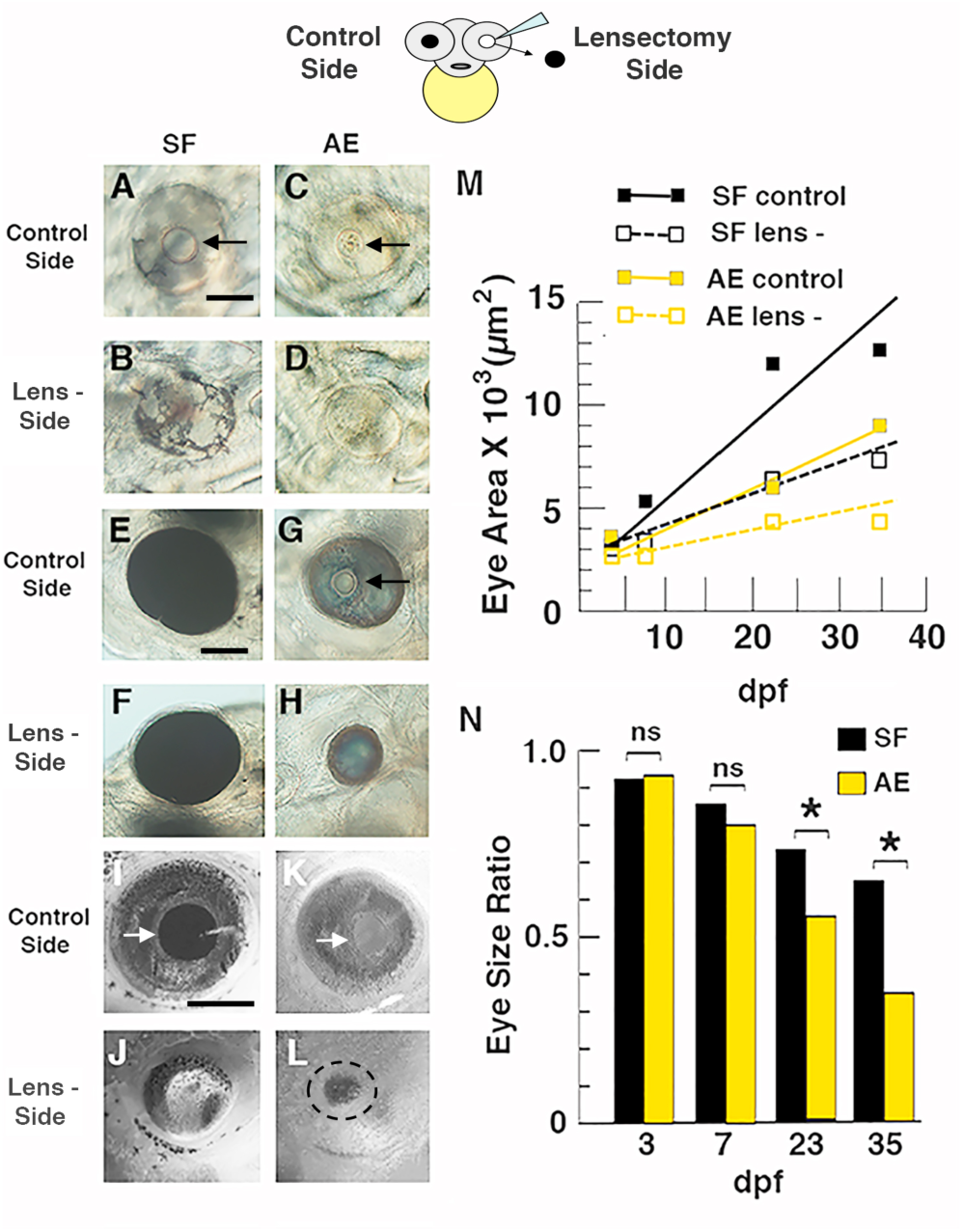
Top. Diagram illustrating the lensectomy experiments shown below and in Figure S2. A-N. Effects of lens deletion on eye size and growth rate in surface fish and F4 AE strain larvae. A-L. Eye size on the lensectomy and control sides of the same larvae at 3 days post-fertilization (dpf) (A-D), 6 dpf (E-H), and 15 dpf (I-L). Arrows: lens. Scale bar in A: 200 μm; magnification is the same in A-D. Scale bar in E: 200 μm; magnification is the same in E-H. Scale bar in I: 300 μm; magnification is the same in I-L. M. Eye growth on the lensectomy and control sides during AE and surface fish development. Best fit lines for eye growth rates were calculated from eye measurements and developmental times using the least squares method. N. Bar graphs showing developmental changes in eye size ratios on the lensectomy and control sides during AE and surface fish development. The data used to illustrate M and N, including means, SEMs, sample numbers, and significance values, are listed in Tables S1 and S2 respectively.

Lens deletion in F4 AE and surface fish larvae resulted in a slight decrease in eyeball size on the operated side relative to the control side at 3 dpf (Fig. 5A-D). The differences in eye ball size were subsequently amplified (Fig. 5 E-H) and became more substantial after several weeks (Fig. 5I-L). By this time, the differences in eyeball size between the operated and control sides of F4 AE were much larger than those of surface fish (Fig. 5G, H, K, L). Almost all of the differences described above were significant (see Table 1 and 2 for F-score and p-values). For example, population (between SF and AE; F_1,92_=163.3), age (3-35 days post fertilization; F_3,92_=184.6) and lensectomy (between control and lensectomy sides, repeated measurement; F_1,92_=304.7) are all highly significant (p < 0.001) (Table 1). In addition, interactions between population and age (F_3,92_=32.7) and between age and lensectomy (F_3,92_=66.3) were also highly significant (p < 0.001)(Table 2). Similar, although smaller, differences were seen in operated F3 AE and corresponding surface fish larvae (Fig. S2A-D), and most of these differences were also significant (Table S1).

Quantification of eyeball size differences was also done by comparing changes in the eye size ratio (defined as the dividend of eyeball size on the operated and control side of the head) during development of F4 AE, F3 AE, and surface fish larvae (Figs. 5N, S2F). The results showed that beginning at 7 dpf eye size ratios between F4 AE and surface fish progressively decreased during larval and juvenile development, and that at 23 dpf (p < 0.0327) and 35 dpf (p < 0.0184) these decreases were significantly larger in F4 AE compared to surface fish larvae (Table 3 and 4). Eye size ratios were also larger in F3 AE relative to surface fish larvae at 10 dpf and 21 dpf (Fig. S2; Table S2), although the differences were significant only at 10 dpf (p < 0.0414). These results suggest that eye growth slows as a function of increasing developmental stage in lens ablated AE compared to surface fish. Furthermore, the results suggest that optic growth on the control side of F3 AE and F4 AE larvae was similar to that on the lens deleted side of surface fish.

In summary, the results of lens deletion experiments indicate that eye development is more sensitive to lens deletion in F3 AE and F4 AE larvae than surface fish larvae, suggesting that the lens and RPE collaborate in promoting the development of full sized eyes in *A. mexicanus*.

### Defective Scleral Differentiation in the Albino Eyed Strain

Previous studies reported evolutionary changes in scleral differentiation in *Astyanax* cavefish: surface fish sclera have a large cartilage ring with two boney ossicles, whereas cavefish develop a much smaller scleral cartilage ring that is generally lacking boney ossicles (Yamamoto et al. 2003, Dufton et al., 2012; O’Quin et al., 2017; Lyon, Powers, Gross, & O’Quin, 2017). To determine the role of the RPE in scleral differentiation, we stained cartilage and bone in F4 AE, surface fish, and cavefish adults of similar age and size. The surface fish and cavefish controls showed the previously described scleral phenotypes (Fig. 6A, D, E). In contrast, large scleral cartilage rings resembling those of surface fish were formed F4 AE adults, but the scleras of the majority of AE adults resembled the cavefish sclera in lacking boney ossicles (Fig. 6B, E). A small number of AE developed one or two boney ossicles (Fig. 6C, E), and some of these were incompletely ossified. The results show that most AE larvae sclera develop large sclera resembling those of surface fish but also resemble cavefish in failing to develop boney ossicles. These results suggest that the RPE has a role in scleral ossification in *A. mexicanus* and is associated with scleral degeneration in cavefish.

**Figure 6.**
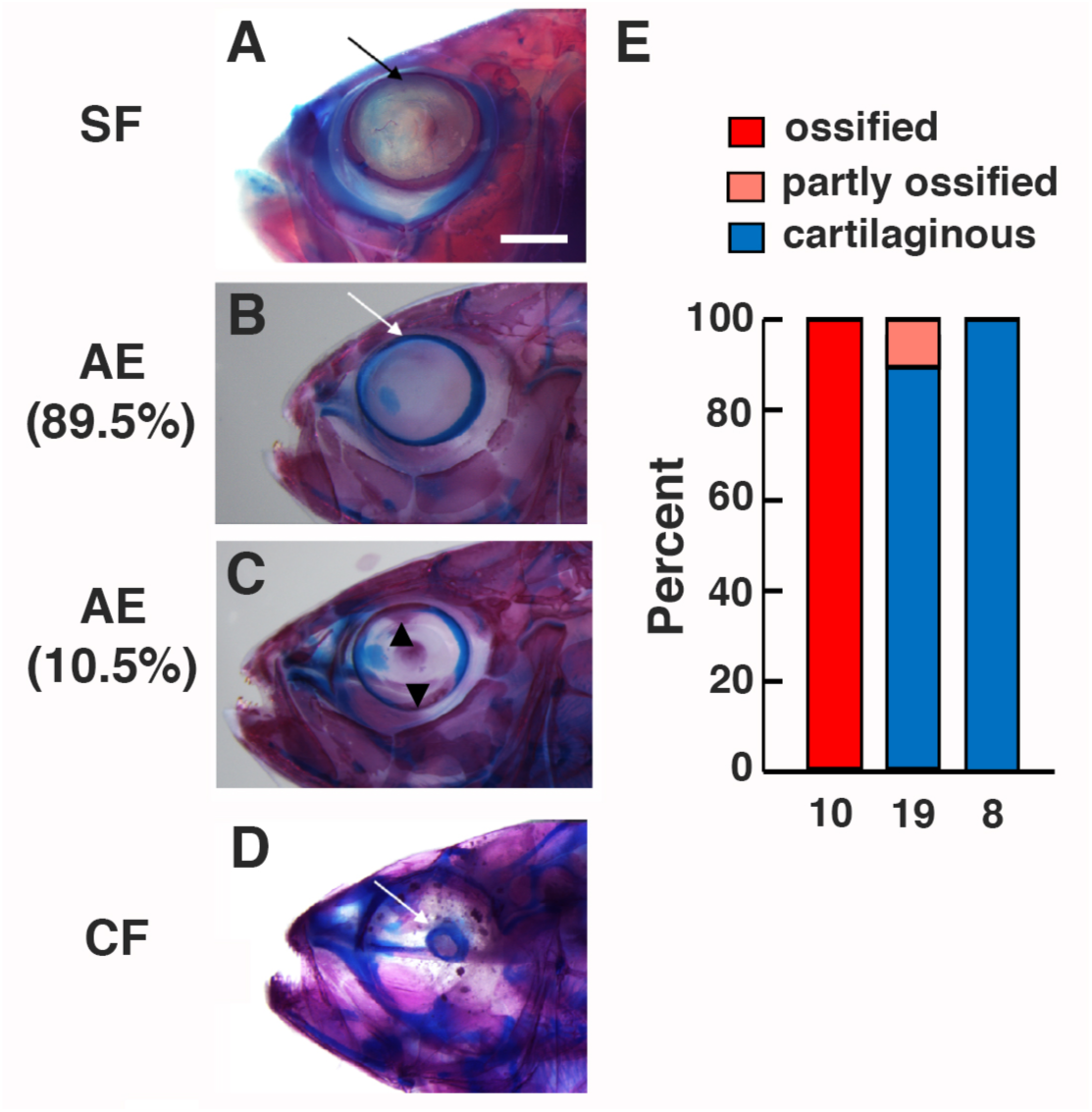
Scleral differentiation in F4 AE strain, surface fish, and cavefish adults. A-D. Surface fish (A), F4 AE strain (B, C), and Pachón cavefish (D) double stained for cartilage in blue and bone in red. Arrows. Scleral rims. Arrowheads in C: Upper and lower scleral ossicles. Percentages under AE in B and C indicate proportion of adults showing scleral traits (see also E). Scale bar in A: 200 μm; magnification is the same in A-D. E. Bar graphs showing the percentage of adults with complete scleral ossification (two fully boney ossicles), partial scleral ossification (one or two fully or partly ossified ossicles), or no scleral ossification in surface fish, F4 AE, and cavefish. N is indicated on the bottom of each bar.

## Discussion

We generated an *A. mexicanus* strain with a defective RPE and large eyes by artificial selection. Comparison of the effects of lens ablation on eye development in the AE strain, which has a defective RPE, and surface fish demonstrated that contributions from both the RPE and lens are required for normal eye growth and suggested that the RPE has an impact on sclera differentiation. Extrapolation of our results to *Astyanax* cavefish suggests that evolutionary changes in development of the RPE and lens were critical factors in the eye regression process.

### Defective Retinal Pigment Epithelium in the Albino Eyed Strain

The results indicate that the defective cavefish RPE phenotype was successfully transferred and merged with the surface fish large eye phenotype in the AE strain by artificial selection. Similar to the RPE of cavefish larvae, melanin pigment granules were absent or rare in the AE RPE, and rpe65 antibody staining showed that retinol isomerase, a key RPE-specific enzyme (Hamel et al., 1993; Jin et al., 2005), is considerably reduced or lacking in AE eyes. Our immunostaining studies also demonstrated that retinal isomerase was present in normal surface fish eyes and not detectable in degenerating cavefish eyes, which have a defective RPE. Together, these findings indicate that the AE strain and cavefish share some of the same deficiencies in RPE development.

The presumptive RPE is formed by a part of the optic vesicle and eventually becomes a monolayer of cells located on the outside of the optic cup and retina during subsequent eye development (Schmitt & Dowling, 1994; Fuhrmann, Zou, & Levine, 2014). Previous studies have demonstrated that the cavefish optic vesicle is reduced in size compared to the surface fish optic vesicle and exhibits changes in the expression boundaries of the critical patterning genes *pax6, pax2*, and *vax1* (Yamamoto, Stock, and Jeffery, 2004). Therefore, it is likely that one of the consequences of modifying the expression territories of optic vesicle patterning genes in cavefish embryos is abnormal RPE development.

Despite exhibiting a defective RPE, the AE strain is similar to surface fish, and differs from cavefish, in developing a relatively normal neural retina. The number and organization of the retinal layers in the AE strain is similar to the surface fish retina, and in most cases the outer layer expresses zpr2 antibody, indicative of the presence of differentiated photoreceptor cells (Qin et al., 2011; Bader et al., 2102). The co-existence of a normal retina and defective RPE in the AE strain appears to conflict with studies implicating a functional and pigmented RPE in proper retinal lamination and photoreceptor cell development in other vertebrates (Raymond & Jackson, 1995; Jeffery, 1997; Longbottom, Fruttiger, Douglas, Martinez-Barbera, Greenwood. et al., 2009; Hanovice et al., 2019). As originally pointed out by Strickler et al. (2007), however, this conflict would be resolved if the lens, which is present in the AE strain, normally functions alone or in concert with the RPE to regulate neuroretinal development. The latter possibility is supported by the rescue of disrupted retinal organization in cavefish eyes that received a transplanted lens from surface fish (Yamamoto & Jeffery, 2000) and recent studies demonstrating that BMP signaling by the lens induces neuroretina development in chick embryos (Patin et al., 2015). Alternatively, the conflicting results could also be explained if the RPE is multifunctional and only the part related to pigmentation, retinol isomerase function, and the control of overall eye growth is altered, whereas another part involved in the organization of retinal development is preserved in the AE strain.

### Multiple Factors Involved in Cavefish Eye Degeneration

Because analysis of mutant phenotypes (Rojas-Muñoz, Dahm & Nüsslein-Volhard, 2005; Gross, Perkins, Amsterdam, Egana, Darland et al., 2005) and genetic ablation (Longbottom et al., 2104; Hanovice et. al., 2019), which have been established to study RPE function in zebrafish, are unavailable in *Astyanax*, we developed an alternative approach using hybrid crosses and artificial selection to study the role of the RPE in eye growth. Accordingly, we tested the requirement of a functional RPE for eye development by comparing the effects of lens ablation between the AE strain, which has a defective RPE, and surface fish, which has a functional RPE. The results showed that lens removal in both the F3 and F4 generations of the AE strain resulted in stronger suppression of eye development, including reduced eye size ratios and optic growth rates, than in surface fish, suggesting that the lens and RPE collaborate to control complete eye development in *A. mexicanus*. Furthermore, these results imply that defects in the both the RPE and lens may be responsible for eye degeneration in *Astyanax* cavefish.

Our results implicating the RPE in cavefish eye degeneration are supported by evidence showing that loss of melanin pigmentation and eye regression are genetically related (Protas et al., 2007). The *Astyanax* QTL map shows several genomic regions in which eye or lens size QTL are coincident, overlapping, or located close to pigmentation QTL (see Protas & Jeffery, 2012 for summary), including the major QTL containing the *oca2* locus, which is responsible for the evolution of albinism in several cavefish lineages (Protas et al., 2006). The clustering of eye and pigmentation QTL could be interpreted as cases of linkage disequilibrium, but they could also reflect the location of genes affecting eye regression through expression pigment cells, either in the RPE, the body, or both. Accordingly, our studies point to a possible explanation for clustering of eye and pigmentation QTL based on collaboration between RPE and/or body pigment cells and the lens in contributing to eye growth in *Astyanax*.

A model summarizing our results based on evidence from the AE strain is shown in Figure 7. According to this model, which is also based in part on previous studies (Yamamoto & Jeffery, 2000; Strickler et al., 2007), signals emanating from both the RPE and lens are required for complete eye development in surface fish (Fig. 7A), and both signals have been lost or modified in cavefish (Fig. 7C), resulting in eye degeneration. The model also proposes that one of the two signals, the putative signal(s) emanating from the RPE, is absent or modified in the AE strain. Furthermore, as described above, functional redundancy between the two signals would result in a normal appearing, albeit smaller retina, in the AE strain because the lens signal would be present but the RPE signal would be modified or missing. The dominant epistatic pattern revealed by quantitative genetic analysis for eye size and scleral ossicle formation (see below) (O’Quin et al, 2013; Lyons et al., 2017) may also be consistent with a dual signal model in that these traits develop normally when both signals are intact but not when both signals are impaired. The identity of the disrupted signals from the lens and RPE remain unknown, but based on studies in other vertebrates there are many reasonable candidates, including BMPs emanating from the lens (Pandit et al., 2015), FGFs (Seidler, Schwegler, & Liesenhoff, 1999), Insulin-like Growth Factors (Waldbillig, Pfeffer, Schoen, Adler, Shen-Orr et al., 1991), or other factors (Strauss, 2005) secreted by the RPE, and Lens Derived Growth Factor produced by both of these eye parts (Nakamura, Singh, Kubo, Chylack, & Shinohara, 2000).

**Figure 7.**
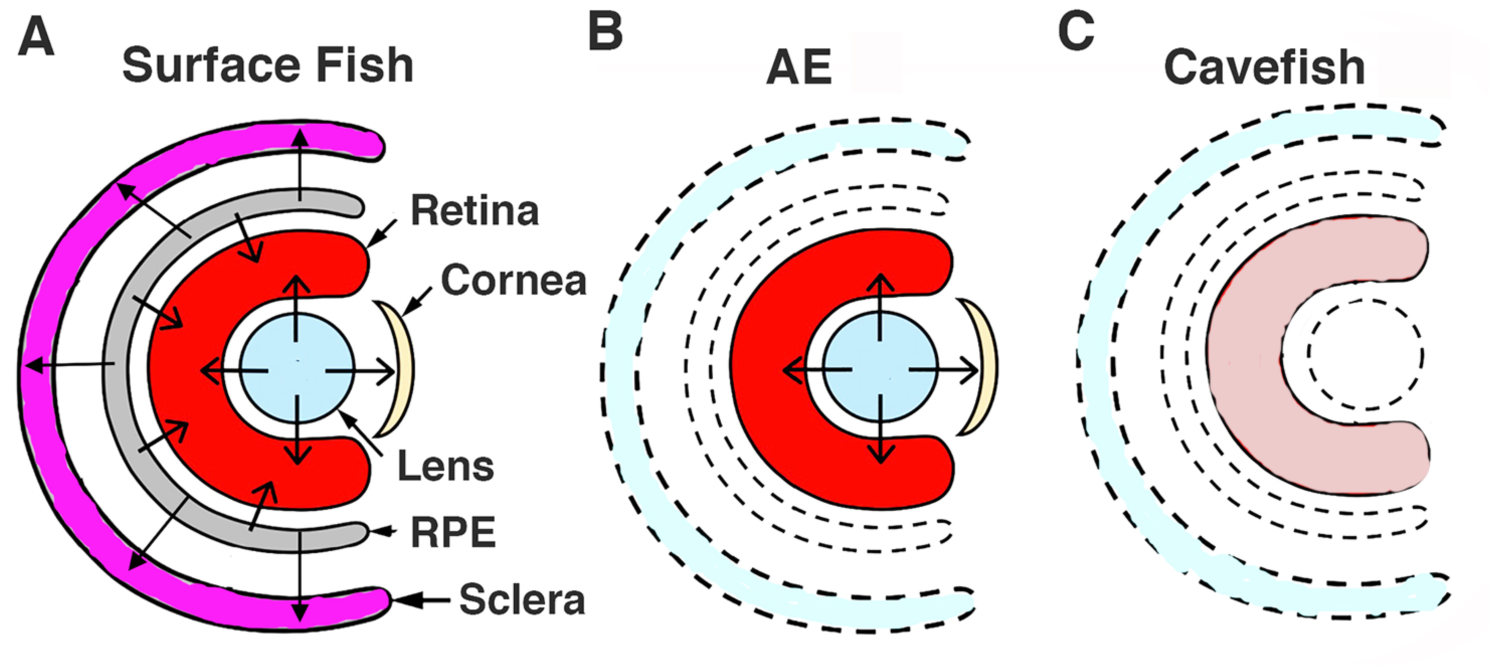
A-C. Model illustrating the hypothesis implicating dual roles of signaling from the lens and RPE in the control of eye development/degeneration in surface fish (A), the albino eyed (AE) strain (B), and cavefish (C). To summarize, in surface fish signaling from the lens and RPE control retinal and sclera development (B), in AE retinal development is normal due to the presence of lens signal but sclera differentiation is compromised due the lack of signaling from the defective RPE, and (C) In cavefish retinal development and scleral differentiation are both compromised due to lack of signaling from the defective lens and RPE. Unlabeled arrows: directions of signals. Drawings do not reflect actual sizes of structures.

Although the model for cavefish eye degeneration (Fig. 7) highlights a dual role of the lens and RPE in cavefish eye degeneration, it is not meant to exclude the possibility that only these two parts of the eye are involved in eye regression. For example, the slightly smaller eyes in AE stain adults could reflect changes in cell numbers or sources of secreted morphogens that may affect eye development independently of the RPE or lens. Likewise, in addition to RPE pigmentation, the absence of body pigmentation outside the eye could affect lens ablation in AE. In recent studies, the optic vasculature, which is ultimately responsible for a healthy choroid layer between the RPE and sclera, has been found to be defective in cavefish (Ma et al., in preparation), and this could also have a major influence on cavefish eye degeneration. Therefore, consistent with genetic evidence showing that multiple genes are involved in the cavefish eye loss (Borowsky &Wilkens, 2001; Protas et al., 2007; O’Quin et al., 2013), it is likely that the physiological basis of cavefish eye degeneration is equally complex, with the involvement of multiple components, including changes in the RPE, lens, and possibly other factors located inside or outside of the eye.

### The RPE Impacts Sclera Ossification

The cavefish sclera is characterized by reduced overall size of the cartilage ring and the absence or reduction of ossicles in some populations (Yamamoto et al., 2003; O’Quin et al., 2015, Lyons et al., 2017). In contrast to the retina, much less is known about the control of sclera development, although there is evidence in chickens and mice suggesting that scleral cartilage formation is dependent on signaling by a functional RPE (Thompson, Griffiths, Jeffery, & McGonnell., 2010). Yamamoto et al. (2003) reported that missing ossified elements of the sclera could be restored by transplantation of a surface fish embryonic lens into the Pachón cavefish optic cup. Furthermore, as is also the case for retinal organization (see above), lens ablation from surface fish did not result in a large reduction in scleral cartilage size or a lack of ossification, as expected if the lens were the only optic tissue responsible for scleral differentiation (Yamamoto et al., 2003; Dufton et al., 2012). These considerations suggest that the dual signal model (Strickler et al., 2007) may also apply to evolutionary changes in cavefish scleral differentiation (Fig. 7).

The demonstration that the F4 AE strain develops a relatively large cartilaginous sclera lacking boney ossicles suggests that a functional RPE is required for scleral ossification. However, in contrast to the results in chick and mouse (Thompson et al., 2010), our results suggest that the *Astyanax* RPE may not be necessary for scleral growth or cartilage differentiation. The qualification pointed out above, namely that only part of a multifunctional RPE may be affected in the AE strain, also applies to this interpretation. Nevertheless, our results add to the current knowledge of sclera development by pinpointing the parts of this structure that may be specifically influenced by the RPE.

It is also noteworthy that the entire sclera (including ossicles, when present) is thought to be derived from the cranial neural crest (Noden, 1983; Hall, 1981), and there is recent evidence that deficiencies in neural crest cell migration may affect scleral development in cavefish (Yoshizawa, Hixon, and Jeffery, 2018). Our studies highlight the complexity of sclera development in *Astyanax*, as well as the multiplicity of control mechanisms responsible for changes in sclera differentiation during cavefish eye degeneration. *Astyanax* cavefish is likely to be a useful model to understand the roles of multiple organizing tissues in sclera development and evolution.

### Artificial Selection and Relationships Between Cavefish and Surface Fish Phenotypes

The present investigation suggests that artificial selection may be a useful tool to understand the relationship between surface fish and cavefish phenotypes if they can be combined in the same hybrid line through artificial selection. Beginning with surface fish x cavefish F2 hybrids, only two rounds of artificial selection were required to generate the AE strain. The limitation of the genetic crossing and artificial selection approach for studying cavefish and surface fish phenotypes requires that the traits under study sort independently in the F2 hybrid generation. Taking this limitation into account, the merger of surface fish and cavefish traits in the same line could be used to confirm the relationship between traits that are suspected to be associated by antagonistic pleiotropy, such as eyes and taste buds or olfactory structures (Yamamoto et al., 2009; Blin et al., 2018). If such traits can be combined into a single line then their complete antagonistic nature would be faulted, whereas if they are not capable of co-existing then a mutually exclusive relationship would be supported. As a proof of principle, the AE stain was recently used to provide evidence supporting a pleiotropic trade-off between melanin pigmentation and catecholamine synthesis in *Astyanax* (Bilandžija et al., 2018).

### Summary and Conclusion

Previous studies suggested a major role of the lens in modulating eye growth (Yamamoto & Jeffery, 2000), but also reported the existence of one or more other determinants of eye development, and it was proposed that one of these elements might be the RPE (Strickler et al., 2007). Our lens ablation results with the AE strain provide evidence that the RPE is involved in cavefish eye degeneration. Therefore, although the lens plays a central role in cavefish eye degeneration, others factors, including the RPE, are also responsible for the evolution of this complex phenotype.

## Acknowledgements

This research was supported by NIH grant EY024941 to WRJ. We thank Craig Foote and Ruby Dessiatoun for maintenance of our *Astyanax* colony, Amy Parkhurst for histology, and Janet Shi for conducting some of the eye measurements.

## References

Alunni, A., Menuet, A., Candal, E., Penigault, J-B., Jeffery, W. R. & Rétaux, S. (2007). Developmental mechanisms for retinal degeneration in the blind cavefish *Astyanax mexicanus*. Journal of Comparative Neurology, 505: 221–233.

Ashery-Padan, R., Marquart, T., Zhou, X., & Gruss, P. (2000). Pax6 activity in the lens primordium is required for lens formation and for correct placement of a single retina in the eye. Genes & Development, 14: 2701–2711.

Atukarala, A. D. S., & Franz-Odendaal, T. A. (2018) Genetic linkage between altered tooth and eye development in lens-ablated *Astyanax mexicanus*. Developmental Biology, 441: 235–241.

Beebe, D. C., & Coates, J. M. (2000). The lens organizes the anterior segment: specification of neural crest cell differentiation in the avian eye. Developmental Biology, 220: 424–431.

Bader, J. R., Kusik, B. W., & Beharse, J. C. (2012). Analysis of KF17 distal tip tracking in zebrafish cone photoreceptors. Vision Research, 75: 37–43.

Bilandžija, H., Ma L, Parkhurst, A., & Jeffery, W. R. (2013). A potential benefit of albinism in *Astyanax* cavefish: Downregulation of the oca2 gene increases L-tyrosine and catecholamine levels as an alternative to melanin synthesis. Public Library of Science ONE, 8 (11): e80823.

Bilandžija, H., Abraham, L., Ma, L, Renner, K., & Jeffery, W. R. (2018). Behavioral changes controlled by catecholaminergic systems explain recurrent loss of pigmentation in cavefish. Proceedings of the Royal Society B, 285: http://doi:10.1098/rspb.2018.0243

Blin, M., Tine, E., Meister, L., Elipot, Y. Bibliowicz, J., Espinasa, L, & Rétaux, S. (2018). Developmental evolution and developmental plasticity of the olfactory epithelium and olfactory skills in Mexican cavefish. Developmental Biology, 441: 242–251.

Borowsky, R., & Wilkens, H. (2002). Mapping a cave fish genome: polygenic systems and regressive evolution. Journal of Heredity, 93, 19–21.

Brakefield P. M., French V., & Zwaan B. J. (2003). Development and the genetics of evolutionary change within insect species. Annual Review of Ecology, Evolution, and Systematics, 34: 633–660.

Breitman, M. L., Clapoff, S, Rossant, J., Tsui, L. C., Glode, M., Maxwell, I. H., & Bernstein, A. (1987). Genetic ablation: targeted expression of a toxic gene causes microphthalmia in transgenic mice. Science, 238: 1563–1565.

Cahn, P. H. (1958). Comparative development in *Astyanax mexicanus* and two of its blind cave derivatives. Bulletin of the American Museum of Natural History, 115: 75–112.

Coulombre, A J., & Coulombre, J. L., (1964). Lens development. I. Role of the lens in eye growth. Journal of Experimental Zoology, 156: 39–48.

Dufton, M., Hall, B. K., & Franz-Odendaal, T. A. (2012). Early lens ablation causes dramatic long-term effects on the shape of bones in the craniofacial skeleton of *Astyanax mexicanus*. Public Library of Science ONE, 7: e50308. pmid:23226260

Fumey, J., Hinaux, H., Noirot, C, Thermes, C., Rétaux, S. & Casane, D. (2018). Evidence for late Pleistocene origin of *Astyanax mexicanus* cavefish. BMC Evolutionary Biology, 18: 18(1):43. doi: 10.1186/s12862-018-1156-7.

Fuhrmann, S., Zou, C-J., & Levine, E. M. (2014). Retinal pigment epithelium development, plasticity, and tissue homoeostasis. Experimental Eye Research, 123: 141–150.

Goldstein, B. (2010). On the evolution of early development in the Nematoda. Philosophical Transactions of the Royal Society London B Biological Sciences, 356: 1521–1531.

Gross, J. B. (2012). The complex origin of *Astyanax mexicanus* cavefish. BMC Evolutionary Biology, 12: 105 http://www.biomedcentral.com/1471-2148/12/105

Gross, J. B., Krutzler, A. J., & Carlson, B. M. 2014. Complex craniofacial changes in blind cave-dwelling fish are mediated by genetically symmetric and asymmetric loci. Genetics, 196: 1303–1319.

Gross, J. M., Perkins, B. D., Amsterdam, A., Egana, A., Darland, T., Matsui, J. I., Sciascia, S., Hopkins, N., & Dowling, J. E. (2005). Identification of zebrafish insertional mutants with defects in visual system function and development. Genetics 170: 245–251.

Hall, B. K. (1981). Specificity in the differentiation and morphogenesis of neural crest-derived scleral ossicles and of epithelial scleral papillae in the eye of the embryonic chick. Journal of Embryology and Experimental Morphology, 66: 175–190.

Hamel, C. P., Tsilou, E., Harris, E., Pfeffer, B. A., Hooks, J. J., Detrick, B., & Redmond, T. M. (1993). A developmentally regulated microsomal protein specific for the pigment epithelium of the vertebrate retina. Journal of Neuroscience Research, 34 (4): 414–25.

Hanovice, N. J., Leach, L. I., Slater, K., Gabriel, A. E., Romanovicz, D, Shao, E, Collery, R., Burton, E. A., Lathrop, K. I., Link, B. A., & Gross, J. M. Regeneration of the zebrafish retinal pigment epithelium after widespread genetic ablation. Public Library of Science Genetic, 15: e1007939. https://doi.org/10.1371/journal.pgen.1007939

Hanken, J., & Wassersug, R. (1981) The visible skeleton. Functional Photography, 16: 22–26.

Herman, A., Brandvain, Y. Weagley, J., Jeffery, W. R., Keene, A. C., Kono, T. J. Y., Bilandžija, H., Borowsky, R., Espinasa, L. O’Quin, K., Ornelas-García, C. P., Yoshizawa, M., Carlson, M., Maldonado, E., Gross, J. B., Cartwright, R. A., Rohner, N., Warren, W. C., & McGaugh, S. E. (2018). The role of gene flow in rapid and repeated evolution of cave related traits in Mexican tetra, *Astyanax mexicanus*. Molecular Ecology, 27: 4397–4416.

Jin, M., Li, S., Moghrabi, W.N., Sun, H., & Travis, G. H. (2005). Rpe65 is the retinoid isomerase in bovine retinal pigment epithelium. Cell, 122: 449–59.

Jeffery, W. R. (2001). Cavefish as a model system in evolutionary developmental biology Developmental Biology, 231: 1–12.

Jeffery, W. R. (2009). Regressive evolution in *Astyanax* cavefish. Annual Review of Genetics, 43: 25–47.

Jeffery, W. R. (2016). The comparative organismal approach in evolutionary developmental biology: insights from ascidians and cavefish. Current Topics in Developmental Biology, 116: 489–500.

Jeffery, W.R., and Martasian, D. P. (1998). Evolution of eye regression in the cavefish *Astyanax*: Apoptosis and the Pax-6 gene. The American Zoologist, 38: 685–696.

Jeffery, W. R., Strickler, A. G., Guiney, S., Heyser, D., & Tomarev, S. I., 2000. Prox1 in eye degeneration and sensory organ compensation during development and evolution of the cavefish *Astyanax*. Development Genes & Evolution, 210: 223–230.

Kaur, S., Key, B., Stick, J., McNeisch, J. D., Akeson, R., & Potter, S. S. (1989). Targeted ablation of alpha-crystallin-synthesizing cells produces lens-deficient eyes in transgenic mice. Development 105: 613–619.

Keene, A. C., Yoshizawa, M., & McGaugh, S. M. (2015). Biology and Evolution of Mexican Cavefish. Elsevier; New York

Klaassen, H., Wang, Y., Adamski, K., Rohner, N., & Kowalko, J. E. (2018). CRISPR mutagenesis confirms the role of *oca2* in melanin pigmentation in *Astyanax mexicanus.* Developmental Biology, 441: 313-318.

Krishnan, J., & Rohner, N. (2017). Cavefish and the basis for eye loss. Philosophical Transactions of the Royal Society London B Biological Science. 5: 372: 20150487. http://dx.doi.org/10.1098/rstb.2015.0487

Kuritia, R., Sagara, H., Aoki, Y., Link, B. A., Arai, K., & Watanabe, S. (2003). Suppression of lens growth by alphaA-crystallin promoter-driven expression of diptheria toxin results in disruption of retinal cell organization in zebrafish. Developmental Biology, 255: 113–127.

Longbottom, R., Fruttiger, M., Douglas, R. H., Martinez-Barbera, J. P., Greenwood, J., & Moss, S. E. (2009). Genetic ablation of retinal pigment epithelial cells reveals the adaptive response of the epithelium and impact on photoreceptors. Proceedings of the National Academy of Sciences of the United States of America, 106: 18728–18733.

Lyon A, Powers AK, Gross JB, & O’Quin KE (2017) Two – three loci control scleral ossicle formation via epistasis in the cavefish *Astyanax mexicanus*. Public Library of Science ONE, 12(2): e0171061. https://doi.org/10.1371/journal.pone.0171061

Ma, L., Strickler, A. G., Parkhurst, A., Yoshizawa, Y., Shi, J., & Jeffery, W. R. (2018). Maternal genetic effects in *Astyanax* cavefish development. Developmental Biology. 441: 209–220.

McCauley, D.W., Hixon, E., & Jeffery, W. R., (2004). Evolution of pigment cell regression in the cavefish *Astyanax*: A late step in melanogenesis. Evolution & Development 6: 209–218.

Nakamura, M., Singh, D. P., Kubo, E., Chylack, L. T. Jr., & Shinohara, T. (2000). LEDGF: survival of embryonic chick photoreceptor cells. Investigative Ophthalmology & Visual Science, 41: 1168–1175.

Noden, D. M. (1983). The role of the neural crest in patterning of the avian skeletal, connective, and muscle tissues. Developmental Biology, 96: 144–165.

O’Quin, K. E., Yoshizawa, M., Doshi, P., & Jeffery, W. R. (2013). Quantitative genetic analysis of retinal degeneration in the blind cavefish. Public Library of Science ONE, 8(2): e57281. doi:10.1371/journal.pone.0057281.

O’Quin, K. E., Doshi, P., Lyon, A., Hoenemeyer, E., Yoshizawa, M., & Jeffery, W. R. (2017). Complex evolutionary and genetic patterns characterize the loss of scleral ossification in the blind cavefish *Astyanax mexicanus*. Public Library of Science ONE, 10(12): e0142208. https://doi.org/10.1371/journal.pone.0142208

Panit, T., Jidigam, V. K., Patthey, C., & Gunhaga, L. 2015. Neural retina identity is specified by lens-derived BMP signals. Development, 142: 1850–1859.

Protas, M. & Jeffery, W. R. (2012). Evolution and development of cave animals: From fish to crustaceans. WIREs Developmental Biology, 1: 823–845

Protas, M.E., Hersey, C., Kochanek, D., Zhou. Y. Wilkens, H., Jeffery, W. R., Zon, L. I., Borowsky, R., & Tabin, C. J. (2006). Genetic analysis of cavefish reveals molecular convergence in the evolution of albinism. Nature Genetics, 38: 107–111.

Protas, M., Conrad, M., Gross, J. B., Tabin, C., & Borowsky, R. (2007). Regressive evolution in the Mexican cave tetra, *Astyanax mexicanus*. Current Biology, 17: 452–454 (2007).

Qin, Z., Kidd III, A. R., Thomas, J. L., Poss, K. S., Hyde, D. R., Raymond, P. A., & Thummel, R. (2011). FGF signaling regulates rod photoreceptor cell maintenance and regeneration in zebrafish. Experimental Eye Research, 93: 726–734.

Raymond, S. M., & Jackson, I. J. (1995). The retinal pigmented epithelium is required for development and maintenance of the mouse neural retina. Current Biology, 5:1286–1295.

Rojas-Muñoz, A., Dahm, R. & Nüsslein-Volhard, C. (2005). *chokh*/*rx3* specifies the retinal pigment epithelium fate independently of eye morphogenesis. Developmental Biology, 288: 348–362.

Romero, A. Green, S. M., Romero, A., Lelonek, M. M., & Stropnicky, K. C. (2003). One eye but no vision: Cavefish with reduced eyes do not respond to light. Journal of Experimental Zoology B. Molecular Evolution Development, 300: 72–79.

Rymer, J., & Wildsoet, C. F. (2005). The role of the retinal pigment epithelium in eye growth regulation and myopia: a review. Visual Neuroscience, 22: 251–261.

Sadoglu, P. (1957). A mendelian gene for albinism in natural cave fish. Experientia, 13: 394–394.

Schmitt, E., & Dowling, J. (1994). Early eye morphogenesis in the zebrafish, *Brachydanio rerio*. Journal of Comparative Neurology, 344: 532–542.

Seidler, D. G., Schwegler, J. S., & Liesenhoff, H. (1999). Expression of fibroblast growth factors 1 and 2 in a human retinal pigment epithelium cell line (K1034). Ophthalmic Research, 31: 280–286.

Strauss, O. 2005. The retinal pigment epithelium in visual function. Physiological Review, 2005: 845–881.

Strickler, A.G., Yamamoto, Y., & Jeffery, W. R. (2001). Early and late changes in *Pax 6* expression accompany eye degeneration during cavefish development. Development Genes & Evolution. 211: 138–144.

Strickler, A. G., Y. Yamamoto, & Jeffery, W. R. (2007). The lens controls cell survival in the retina: evidence from the blind cavefish *Astyanax*. Developmental Biology, 311: 512–523.

Strickler, A.G., & Jeffery, W. R. (2009). Differentially expressed genes identified by cross species microarray in the blind cavefish *Astyanax*. Integrative Zoology, 4: 31–40.

Teyke, T. (1990). Morphological differences in neuromasts of the blind cave fish *Astyanax hubbsi* and the sighted river fish *Astyanax mexicanus*. Brain Behavior and Evolution, 35: 23–30

Thompson, H, Griffiths, J. S., Jeffery, G., & McGonnell, I. M. (2010). The retinal pigment epithelium of the eye regulates the development of scleral cartilage. Developmental Biology, 347: 40–52

Yamamoto, Y., & Jeffery, W. R. (2000). Central role for the lens in cave fish eye degeneration. Science, 289: 631–633.

Yamamoto, Y., & Jeffery, W. R. (2002). Probing vertebrate eye development by lens transplantation. Methods, 28: 420–426.

Yamamoto, Y., Espinasa, L, Stock, D. W., & Jeffery, W. R. (2003). Development and evolution of craniofacial patterning is mediated by eye-dependent and -independent processes in the cavefish *Astyanax*. Evolution & Development, 5: 435–446.

Yamamoto, Y., Stock, D. W., & Jeffery, W. R. (2004). Hedgehog signalling controls eye degeneration in blind cavefish. Nature, 431: 844–847.

Yamamoto, Y., Byerly, M. S., Jackman, W. R., & Jeffery, W. R. (2009). Pleiotropic functions of embryonic sonic hedgehog expression link jaw and taste bud amplification with eye loss during cavefish evolution. Developmental Biology, 330: 200–211.

Yoshizawa, M. (2015). Behaviors of cavefish offer insight into developmental evolution. Molecular Reproduction & Development, 82: 268–280.

Yoshizawa, M., Gorički, Š., Soares, D., & Jeffery, W. R. (2010). Evolution of a behavioral shift mediated by superficial neuromasts helps cavefish find food in darkness. Current Biology, 20: 1631–1636.

Yoshizawa, M., Yamamoto, Y., O’ Quin, K. E., & Jeffery, W. R. (2012). Evolution of an adaptive behavior and its sensory receptors promotes eye regression in blind cavefish. BMC-Biology, 10:108 doi:10.1186/1741-7007-10-108.

Yoshizawa, M., Jeffery, W. R., van Netten, S. M, and McHenry, M. J. (2014). The sensitivity of lateral line receptors and their role in behavior of Mexican blind cavefish (*Astyanax mexicanus*). Journal of Experimental Biology, 217: 886–895.

Yoshizawa, M. Hixon, E., & Jeffery, W. R. (2018). Neural crest transplantation reveals key roles in the evolution of cavefish development. Integrative & Comparative Biology, icy006, https://doi.org/10.1093/icb/icy006

Waldbillig, R. J., Pfeffer, B. A., Schoen, T. J., Adler, A. A., Shen-Orr, Z., Scavo, L., Le Roith, D. E., & Chader, G. J. (1991). Evidence for an insulin-like growth factor autocrine-paracrine system in the retinal photoreceptor-pigment epithelial cell complex. Journal of Neurochemistry, 57: 1522–1533.

Wilkens, H., (1988). Evolution and genetics of epigean and cave *Astyanax fasciatus* (Charicidae, Pisces). Evolutionary Biology, 23: 271–367.

Zilles, K., Tillmann, B., & Bennemann, R. (1983). The development of the eye in *Astyanax mexicanus* (Caharacidae, Pisces), and its blind cave derivative *Anoptichthys jordani* (Characidae, Pisces), and their crossbreeds. A scanning and transmission electron microscopic study. Cell and Tissue Research, 229: 423–432.

