## Supplementary Files for "Dual Roles of the Retinal Pigment Epithelium and Lens in Cavefish Eye Degeneration"

**Supplementary Information**

**Supplementary Figures and Legends**

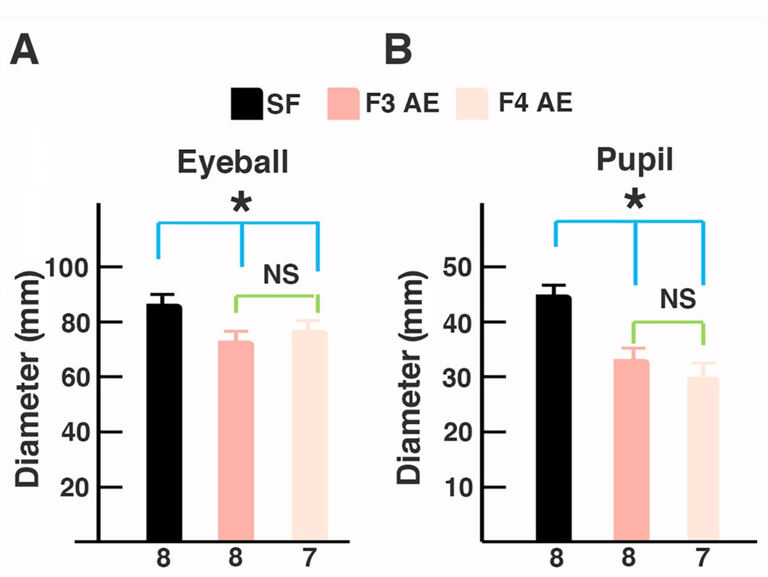


Figure S1. Eye and pupil sizes in the albino eyed (AE) strain. A, B. Bar graphs showing eyeball (A) and pupil (B) diameters in adult F3 AE and F4 AE compared to surface fish of the same size and age. Asterisks indicate significant differences of $p = 0.000$. Error bars: SEM. NS: no significant differences. Statistical analysis by one-tailed Student's t test.

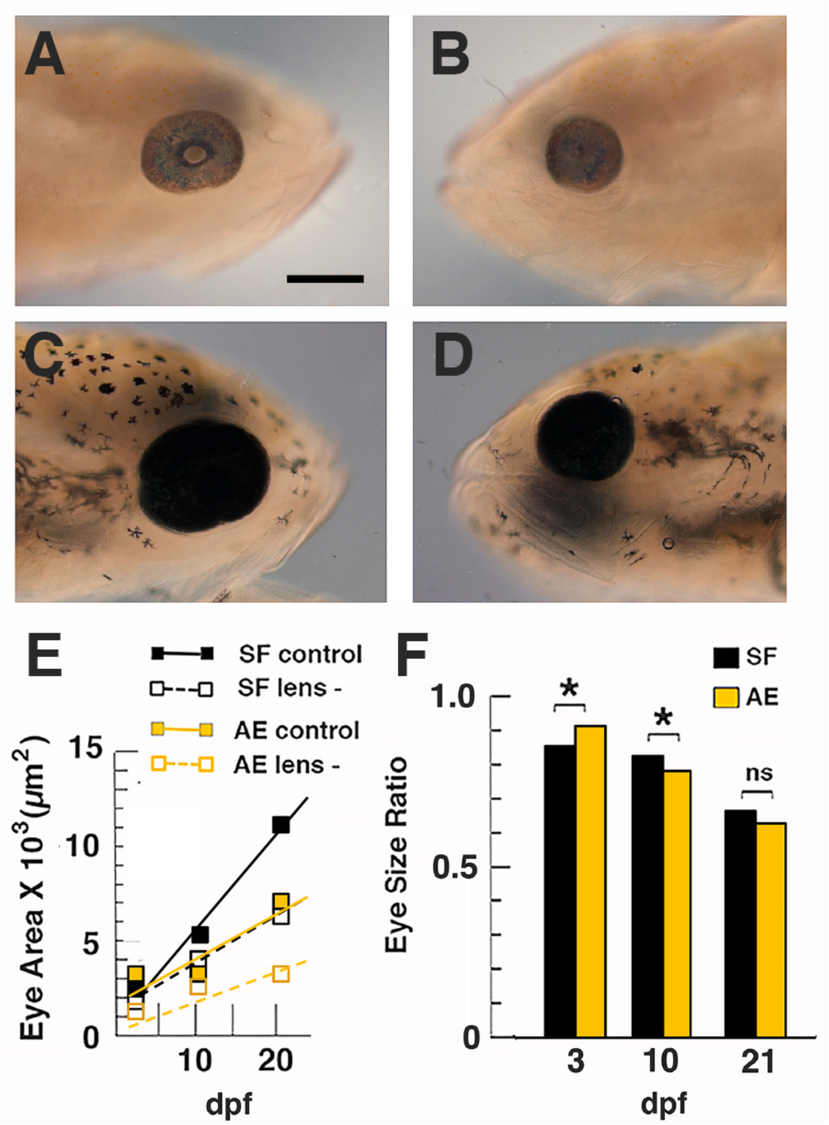


Figure S2. Effects of lens deletion on eye size and growth rate in surface fish and F3 AE strain larvae. A-D. Eye size on lensectomy side and control side of the same surface fish (A, B) and AE (C, D) larvae at 10 days post fertilization (dpf) (I-L). Scale bar in A: 300 $\mu$m; magnification is the same in A-D. E. Eye growth rate on the lensectomy and control sides during AE and surface fish development. F. Bar graphs showing developmental changes in the eye size ratios on the lensectomy and control sides during surface fish and SE development. The data used to prepare E and F, including means, SEMs, sample numbers, and significance values, are listed in Tables S3 and S4 respectively. Other details are the same as in Figure 5.

**Supplementary Tables**

Table S1. Effect of Lens Deletion on Eye Size in Developing Progeny of F3 Albino Eyed (AE) Strain and Surface Fish (SF)

| Age (dpf) † | Type | Pooled Number | Mean Control Side $\mu m^2$ +/- SEM | Mean Lensectomy Side $\mu m^2$ +/- SEM | F¶ | p¶ |
|---|---|---|---|---|---|---|
| 3 | SF | 6 | 253.2 +/- 7.9 | 216.3 +/- 7.6 | 11.3319 | 0.0281 |
| 3 | AE | 8 | 251.4 +/- 5.4 | 234.3 +/- 3.8 | 6.5665 | 0.0428 |
| 10 | SF | 6 | 511.0 +/- 13.2 | 426.6 +/- 3.0 | 38.5149 | 0.0034 |
| 10 | AE | 7 | 448.2 +/- 12.2 | 325.3 +/- 17.5 | 28.6374 | 0.0031 |
| 21 | SF | 11 | 1153.1 +/- 39.1 | 804.5+/- 43.53 | 34.0891 | 0.0002 |
| 21 | AE | 8 | 727.9 +/- 91.0 | 491.9 +/- 102.7 | 2.9781 | 0.1352 |

†dpf: Days post fertilization
‡Significance determined by One-way ANOVA with post-hoc Bonferroni correction.

Table S2. Eye Size Ratio of Lensectomy and Control Eyes in Developing Progeny of F3 Albino Eyed Strain (AE) and Surface Fish (SF)

| Age (dpf) † | Type‡ | Eye Size Ratio§+/- SEM | F¶ | p¶ |
|---|---|---|---|---|
| 3 | SF | 0.867 +/- 0.017 | 21.0815 | 0.0059 |
| 3 | AE | 0.938 +/- 0.059 | | |
| 10 | SF | 0.836 +/- 0.021 | 8.7950 | 0.0414 |
| 10 | AE | 0.767 +/- 0.802 | | |
| 21 | SF | 0.691 +/- 0.034 | 0.0930 | 0.8690 |
| 21 | AE | 0.670+/- 0.131 | | |

†dpf: Days post fertilization
‡Number of specimens is same as in Table S3.
§Ratio is mean eye size on lensectomy side/control side.
¶F-statistic and p value significance between SF and AE determined by One-way ANOVA with post-hoc Bonferroni correction.
